# Calculating functional diversity metrics using neighbor-joining trees

**DOI:** 10.1101/2022.11.27.518065

**Authors:** Pedro Cardoso, Thomas Guillerme, Stefano Mammola, Thomas J. Matthews, Francois Rigal, Caio Graco-Roza, Gunilla Stahls, Jose Carlos Carvalho

**Affiliations:** Laboratory for Integrative Biodiversity Research (LIBRe), Finnish Museum of Natural History (Luomus), University of Helsinki, Helsinki, Finland; CE3C—Centre for Ecology, Evolution and Environmental Changes/Azorean Biodiversity Group / CHANGE – Global Change and Sustainability Institute and Universidade dos Açores – Faculty of Agricultural Sciences and Environment, Angra do Heroísmo, Açores, Portugal; Department of Animal and Plant Sciences, The University of Sheffield, Sheffield, UK; Molecular Ecology Group (MEG), Water Research Institute, National Research Council (CNR-IRSA), Verbania Pallanza, Italy; GEES (School of Geography, Earth and Environmental Sciences) and Birmingham Institute of Forest Research, University of Birmingham, Birmingham, UK; Université de Pau et des Pays de l’Adour, CNRS, Pau, France; Department of Geosciences and Geography, University of Helsinki, Helsinki, Finland; Laboratory of Ecology and Physiology of Phytoplankton, Department of Plant Biology, State University of Rio de Janeiro, Rio de Janeiro, RJ, Brazil; Finnish Museum of Natural History (Luomus), University of Helsinki, Helsinki, Finland; Molecular and Environmental Centre - Centre of Molecular and Environmental Biology, Department of Biology, University of Minho, Braga, Portugal

**Keywords:** Convex hulls, dendrograms, functional divergence, functional diversity, functional regularity, functional traits, hypervolumes, neighbor-joining

## Abstract

1. The study of functional diversity (FD) provides ways to understand phenomena as complex as community assembly or the dynamics of biodiversity change under multiple pressures. Different frameworks are used to quantify FD, either based on dissimilarity matrices (e.g., Rao entropy, functional dendrograms) or multidimensional spaces (e.g. convex hulls, kernel-density hypervolumes). While the first does not enable the measurement of FD within a richness/divergence/regularity framework, or results in the distortion of the functional space, the latter does not allow for comparisons with phylogenetic diversity (PD) measures and can be extremely sensitive to outliers.
2. We propose the use of neighbor-joining trees (NJ) to represent and quantify functional diversity in a way that combines the strengths of current FD frameworks without many of their weaknesses. Our proposal is also uniquely suited for studies that compare FD with PD, as both share the use of trees (NJ or others) and the same mathematical principles.
3. We test the ability of this novel framework to represent the initial functional distances between species with minimal functional space distortion and sensitivity to outliers. The results using NJ are compared with conventional functional dendrograms, convex hulls, and kernel-density hypervolumes using both simulated and empirical datasets.
4. Using NJ we demonstrate that it is possible to combine much of the flexibility provided by multidimensional spaces with the simplicity of tree-based representations. Moreover, the method is directly comparable with PD measures, and enables quantification of the richness, divergence and regularity of the functional space.

## Introduction

With the advent of new ways of thinking about biodiversity (McGill et al., 2006) and novel sources of data (Jarić et al., 2020; Tosa et al., 2021; Tobias et al., 2022), we are experiencing a shift from measuring biodiversity based on species identities only (taxonomic diversity, TD), to taking into account either species evolutionary relationships (phylogenetic diversity, PD) or similarities in functional traits (functional diversity, FD) (Pavoine & Bonsall, 2011). An integrative approach to quantifying biodiversity enables not only the comparison of its multiple facets (Pollock et al., 2020), but provides new tools to understand phenomena as complex as community assembly or the dynamics of biodiversity change under multiple pressures (McGill et al., 2006).

Both PD and FD can be measured within (alpha diversity) and between samples, sites or time steps (beta diversity) (Whittaker, 1960). Diversity can also be measured in terms of richness, divergence and regularity (Pavoine & Bonsall, 2011; Tucker et al. 2017; Mammola et al. 2021). For PD, these facets are usually quantified using dissimilarity matrices, either directly from the raw dissimilarity (e.g., Rao entropy, Botta-Dukat 2005; Hill numbers, Chao et al. 2014) or from phylogenetic trees (Tucker et al., 2017). For FD, multidimensional approaches reflecting the niche concept by Hutchinson (1957) are often used, with multiple advantages (Blonder, 2016; Carvalho & Cardoso, 2020; Mammola & Cardoso, 2020; Mammola et al., 2021) over tree-based metrics (Petchey & Gaston, 2002, 2006). For example, functional trees are usually built using hierarchical clustering methods such as the Unweighted Pair Group Method using arithmetic Averages (UPGMA; Michener & Sokal, 1957; Cardoso et al., 2014) or single-linkage trees (equivalent to minimum spanning trees; Villeger et al., 2008), and it has been shown that, in comparison, hyperdimensional approaches better maintain the original distances between species, which eliminates or at least minimises the distortion of the functional space (Maire et al., 2015).

Trees and hyperdimensional representations however require the use of different methods with non-comparable mathematical properties (Mammola et al., 2021). This way, when comparing phylogenetic and functional diversity, it is impossible to know if any differences observed in index values between samples (e.g., richness) are due to inherent differences of the studied systems or due to the use of different algorithms. Using dissimilarity matrices or trees is the only option to compare PD and FD. As many studies use phylogenetic trees, and many regularity and beta diversity partitioning metrics are exclusively calculated using functional tree representations, the use of trees is often preferred for PD/FD comparisons (Cardoso et al., 2014).

Phylogenetic tree reconstruction has seen major advances in recent decades due to the development of ever more complex and efficient algorithms for the representation of evolutionary relationships (Nguyen et al., 2015; Minh et al., 2020). Among the most used, the Neighbor-Joining (NJ) method reconstructs (phylogenetic) trees from evolutionary distance data (Saitou & Nei, 1987). The algorithm for constructing NJ trees connects the terminals based on their overall similarity, and continues to be widely used as it is known to be both efficient and computationally fast. It has a much better performance for reconstructing distance-based trees than UPGMA (Saitou & Nei, 1987). Even if other methods can outperform it under different evolutionary scenarios (e.g., Rannala & Yang, 1996), NJ trees are still widely used for preliminary similarity clustering at the species level. As an example, the Taxon ID tree tool in BOLD (Ratnasingham & Hebert, 2007) employs NJ under the Kimura two-parameter (K2P; Kimura, 1980) distance metric. At the species level, NJ performs best when the mutational rate heterogeneity among the terminals is low, and typically resolves a well-sampled input distance matrix consistently (e.g., Atteson, 1997). The tree topology is also correct if each entry in the distance matrix differs from the true distance by less than half of the shortest branch length in the tree (Mihaescu et al., 2009). As both NJ and UPGMA lack an optimality criterion defining the best tree, an analysis returns only one optimal topology. Finally, all diversity measures calculated using trees constructed using alternative methods can also be calculated using NJ trees with no changes required, this way allowing comparison of, for example, phylogenetic trees constructed using Bayesian methods with functional NJ trees.

In this work, we propose and test NJ trees as a way to quantify richness, divergence and regularity of FD without the functional space distortion typical of functional dendrograms built using hierarchical clustering. We propose a new framework to quantify different dimensions of FD using trees constructed with the Neighbor-Joining algorithm, although the same measures can be applied to trees constructed with any algorithm, from hierarchical clustering to maximum parsimony or maximum likelihood, thus enabling straightforward comparisons of FD with PD. We provide functions for all methods in the R package BAT (Cardoso et al., 2015).

## Materials and methods

Building NJ trees requires a distance matrix between pairs of taxa. In the construction of phylogenetic trees using NJ, the principle of parsimony is used. The algorithm builds a non-ultrametric tree, in a way that the branching patterns and branch lengths are optimised to minimise the amount of change needed to connect all species along the tree (Saitou & Nei, 1987). Hence, the total length of the tree, equivalent to a measure of phylogenetic richness, is also minimised (Fig. 1). Here we propose to use the same principle to build functional trees depicting distances between species (function *BAT::tree.build*). For taxonomic diversity, a star-like tree could be used, with all pairwise distances being equal to one, this way guaranteeing comparability for TD, PD and FD.

**Fig. 1.**
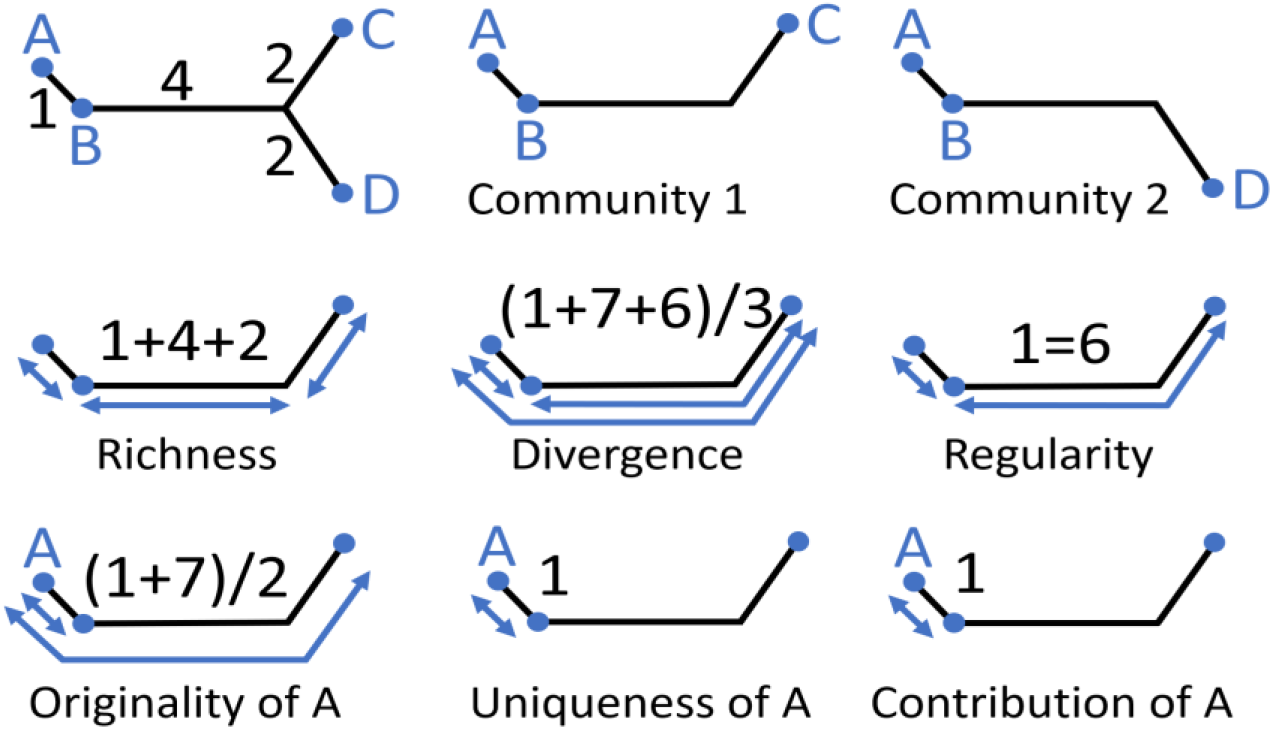
Top-left: NJ tree representing evolutionary or functional distances (edges, in black) connecting four hypothetical species A to D (nodes, in blue). A:B = 1; A:C = A:D = 7; B:C = B:D = 6; C:D = 4. Top-centre and top-right: the same tree for two communities with species A, B and C, and A, B and D respectively. In the middle we represent the calculation of different metrics for community 1. At the bottom, different metrics for species A within community 1 (see main text).

In the context of FD, the same algorithm is used but replacing phylogenetic distance with functional distance between the species. Similar to the concept of hierarchical clustering or minimum spanning trees (which are in effect equivalent to hierarchical clustering with single linkage; Gower & Ross, 1969), NJ trees are much more flexible due to two characteristics. First, they do not explicitly build ultrametric trees as in UPGMA and related methods. Second, as NJ resolves nodes dichotomously, intermediate nodes are introduced in the tree, not forcing species to be in such nodes, as with minimum spanning trees. Such flexibility results in cophenetic distances much closer to the initial distances, decreasing the known limitations of hierarchical clustering that distorts this relation (Maire et al., 2015).

When calculating FD, the distances between species can be Euclidean if all traits are continuous, and traits should generally be standardised (e.g., z-score) to ensure the weight of each is similar (*BAT::standard*). As categorical or ordinal traits are commonly available, the Gower distance is often used (Pavoine et al., 2009; *BAT::gower*), and it is possible to weight traits if some are considered more important with regard to how species interact with their environment, including other species. As traits are often correlated, one might want to eliminate putative relations by first performing a PCoA and using the resulting orthogonal axes as the new traits.

For all analyses one should always use the same phylogenetic or functional tree depicting the relationships between all species, to guarantee comparability of results. Phylogenetic or functional richness is the sum of lengths of edges connecting all species in a community (Faith, 1992; Petchey & Gaston, 2002, 2006; *BAT::alpha*). For the communities in Fig. 1 richness would be Community 1 = Community 2 = 7. Species-level measures can be calculated in different ways. Originality of a species is measured as its average distance to all other species in a community (Pavoine et al., 2005; *BAT::originality*). In the example, for Community 1 the originality would be A = 4, B = 3.5 and C = 6.5. Uniqueness of a species is measured as its distance to the single closest species in the community (Mouillot et al., 2013; *BAT::uniqueness*). In Community 1 it would be A = 1, B = 1, and C = 6. The contribution of a species to richness or alpha diversity is the length of branches unique to it, plus the proportional length of shared branches connecting it to the root of the tree (Isaac et al., 2007; *BAT::contribution*). As NJ trees are unrooted, they can be rooted at the species with minimum originality, which is the one closer to the centre of the NJ tree. In Community 1, contribution would be A = 1, B = 0, and C = 6. This option for rooting is however mostly arbitrary, and alternatives could be explored in the future.

The second dimension of phylogenetic or functional alpha diversity is divergence (Mammola et al., 2021). It can be calculated as the average dissimilarity between any two species or individuals randomly chosen in a community (*BAT::dispersion*). If abundance data are used, dispersion is the quadratic entropy of Rao (1982), otherwise it is the phylogenetic dispersion measure of Webb et al. (2002). In the example of Fig. 1, if all species abundances are 1, dispersion would be Community 1 = Community 2 = 4.667.

The third dimension of phylogenetic or functional alpha diversity is regularity (Mammola et al., 2021). It represents the evenness in the abundances and distances between connected species in a community (*BAT::evenness*). It can be calculated, among others, based on the index of Camargo (1993). In the example of Fig. 1, if all species abundances are 1, evenness would be Community 1 = Community 2 = 0.754.

Finally, beta diversity represents the dissimilarity between two communities (*BAT::beta;* measured using either Jaccard or Soerensen dissimilarity) and can be partitioned into the two processes contributing to it, replacement and loss or gain of species leading to differences in richness (Carvalho et al., 2012), evolutionary history, or functional traits (Cardoso et al., 2014). In the example, comparing communities 1 and 2, β_total_ =β_repl_ = 0.444, and β_rich_ = 0.

### Comparing frameworks using simulated scenarios

We simulated trees using a birth-death model, with both birth (speciation) and death (extinction) parameters drawn from a uniform distribution (0, 1) while keeping the death parameter lower than the birth parameter. For each lineage simulated in the birth-death process, we also simultaneously simulated a trait value as a function of branch length. The traits were simulated using either: 1) a Brownian motion process (whereby the trait value at time t+1 is independent of its value at time t, resulting in an increase of trait variance through time); or 2) an Ornstein-Uhlenbeck process (whereby the trait value at time t+1 is independent of its value at time t but constrained by an overall parameter alpha effectively reducing the increase of trait variance through time). For each tree simulation, we chose the trait process randomly between both processes described above. We ran the birth-death and trait simulations until reaching 100 co-occurring species. For each simulation, we then discarded the extinct species resulting in trees with 100 tips with 1 trait value each. We ran the birth-death and trait simulations using the R package *dads* (Guillerme, 2022).

For each of the two evolutionary processes, we simulated 1, 2, 4, or 8 orthogonal (i.e., uncorrelated) traits, 10 runs per trait number combination, totaling 40 runs per process. For each of these 40 runs the simulation created a functional tree for 100 extant species. For each run, we then sampled 10 communities with an increasing number of species (10, 20,…, 100 species), reaching a final sample size of 800 (2 processes * 4 sets of number of traits * 10 runs * 10 communities).

For each community, we first estimated the correlation between functional richness, divergence, and regularity calculated with NJ trees, checking whether the three metrics were able to capture distinct facets of FD (which is achieved when correlation is low; Mouchet et al., 2010). Next, we used the BAT R package to estimate and compare functional richness, divergence, and regularity with NJ trees, UPGMA trees, and kernel-density n-dimensional hypervolumes. We used Spearman’s correlation to compare the frameworks.

### Functional space quality and sensitivity to outliers

When building a functional space, a crucial aspect is to assess its quality, i.e., the extent to which the functional space is an accurate representation of the initial trait values. In order to achieve this goal, for each pair of species i and j, we compared the initial dissimilarity distance (d_ij_) with the distance in the functional space (h_ij_) obtained by NJ, UPGMA and PCoA (multidimensional space) methods. For the NJ and UPGMA trees, h_ij_ corresponded to the cophenetic distance between species i and j. As for PCoA, we calculated the Euclidean distance between the coordinates of species i and j in the space defined by the PCoA axes. We then calculated the quality of the representation of the functional spaces using the same three frameworks (NJ, UPGMA and PCoA) using the functions *BAT::tree.quality* and *BAT::hyper.quality* (the latter being used for any representation using hyperspaces, i.e., convex hulls or kernel-density hypervolumes). Both these functions calculate the inverse of mean squared deviation between initial and cophenetic distances (Maire et al., 2015) after standardisation of all values between 0 and 1 for simplicity of interpretation and comparability of trees and multidimensional spaces.

The quality of the functional spaces was evaluated in 10 simulations for each combination of number of species per community (from 10 to 100 species), number of traits per species (one, two, four and eight) and evolutionary processes used to generate the traits (BM and OU). For PCoA we did not assess the quality for single traits. It is worth noting that the maximum number of PCoA axes that can be extracted from a matrix of N continuous traits is N. Hence, the quality of the functional space is 1 when using N axes. Therefore, we only used the simulated datasets with eight traits to assess the quality of the functional spaces built by PCoA.

We used linear mixed models to estimate the effect of the different methods (fixed effects) in the quality of the functional space. The number of species per community, the number of traits per species and the evolutionary process used to generate the traits were introduced in the models as random effects. Because the quality of the functional space ranges between 0 and 1, with true 1s but no true 0s included in the response, we transformed all 1s by subtracting 0.0001 and then ran the models following a beta distribution. Mixed models were performed using the glmmTMB (Mollie et al., 2017) package, and model validation was performed by checking heterocedasticity, posterior predictive checking, and normality of random effects and residuals using the performance (Lüdecke et al., 2021) package in R (R Core Team, 2021).

Finally, the sensitivity to outliers was also compared by deleting the species with higher uniqueness in each community and calculating the percentage of change in the values of richness before and after deletion. For the multidimensional space, we calculated differences in richness for kernel-density hypervolumes.

### Comparing frameworks using empirical data

The study of avian functional diversity has recently gained momentum due to the release of the AVONET database, which provides a complete set of data for eight continuous morphological traits for all the world’s extant bird species (Tobias et al., 2022). Dozens of papers have been published using this data source in just a few months (e.g., Weeks et al., 2022), including several focusing on islands (e.g., Matthews et al., 2022; Soares et al., 2022). An often mentioned issue when studying the functional diversity of birds is the so-called “kiwi problem”. In short, kiwis (Apterygidae: *Apteryx* spp.) differ substantially from other birds regarding their morphology (e.g., for wing length, in AVONET, the kiwis have values roughly 267 times smaller than the species with the next smallest wing length) and thus all five species are always (extreme) outliers in functional diversity analyses. As such, there can be large differences in (functional) richness depending on whether they are included when building the functional space or not (e.g., Matthews et al., 2022; see also Fig. 1 in Pigot et al., 2020). These differences reflect the sensitivity to outliers of multidimensional representations such as convex hulls and kernel-density hypervolumes.

To test the sensitivity to outliers of UPGMA, NJ, convex hulls and kernel-density n-dimensional hypervolumes, we took the five kiwi species and then randomly selected 100 bird species from the global species pool, making sure to include one representative from each order. For these 105 species, we sourced data on eight continuous traits (total beak length from the tip to the skull, beak length to the nares, beak width and depth at the nares, wing length, secondary length, tail length, and tarsus length) from AVONET (Tobias et al., 2022). Traits were log-transformed and scaled to mean = 0, sd = 1. We then built the UPGMA and NJ trees from these data after calculating Euclidean distances between species. For both convex hulls and kernel hypervolumes we selected the first five axes of a PCA that summed to 99% explained variance to avoid the use of correlated variables. Functional richness was calculated with the *BAT::alpha, BAT::hull.alpha* and *BAT::kernel.alpha* functions. We then re-calculated functional richness after removing the kiwis from the community and quantified the percent loss. In addition, we quantified the tree and hyperspace qualities as above.

## Results

In regard to the simulations, the correlation was low for all trait combinations, except for richness versus regularity which attained values around 0.7 or above (Fig. 2). Richness and regularity were also sensitive to the number of species (r > 0.6 for all trait combinations), as expected, at least for richness. We found a very high convergence among the estimations based on NJ and UPGMA trees, irrespective of the number of traits and the facets of FD (Fig. 3). Correlations were lower between NJ and hypervolumes, especially for the divergence and regularity components, but also for the richness component in high dimensions (eight traits).

**Fig. 2.**
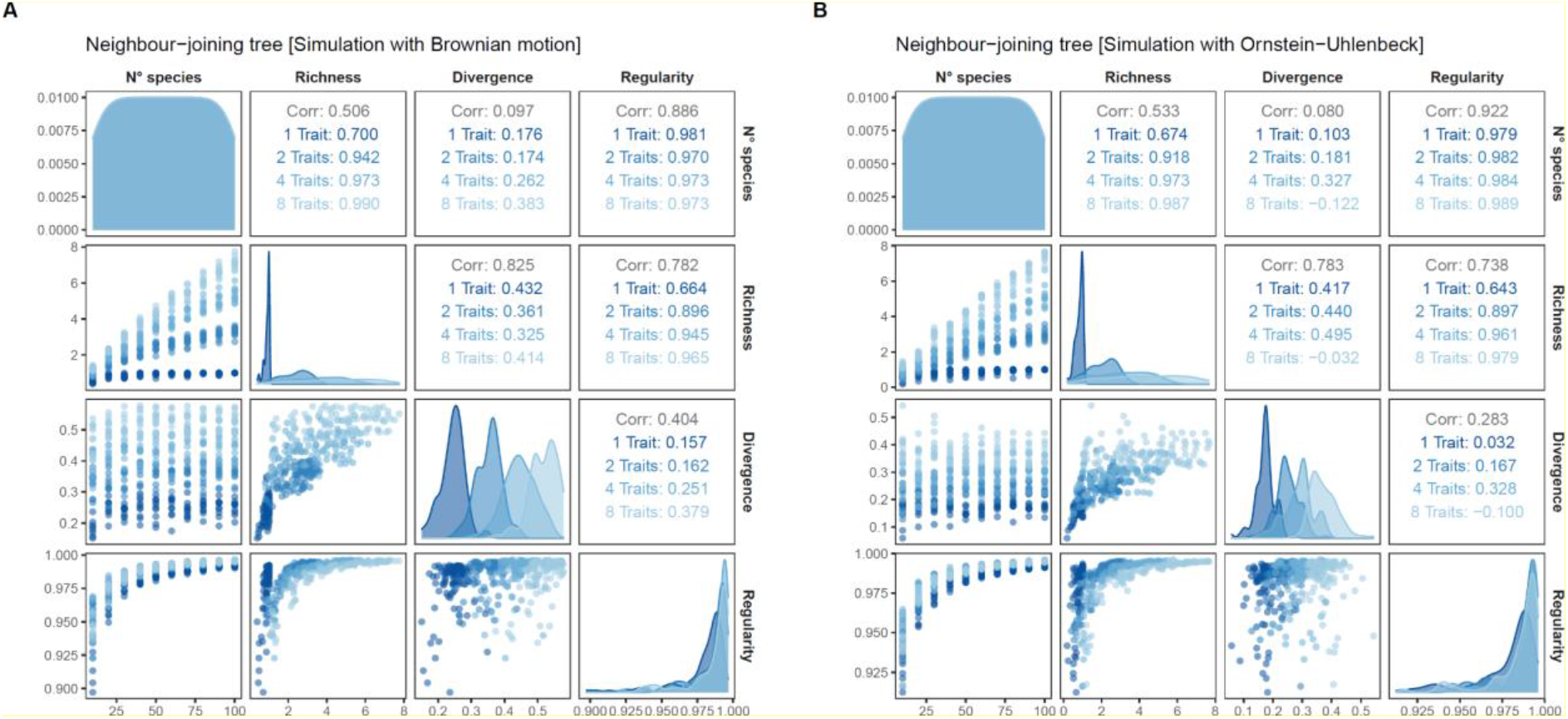
Pairwise Spearman’s correlations among estimations of functional richness, divergence, and regularity based on NJ trees. Density plots on the diagonal display the distribution of values. Bivariate scatter plots are displayed below the diagonal and the correlation values above the diagonal.

**Fig. 3.**
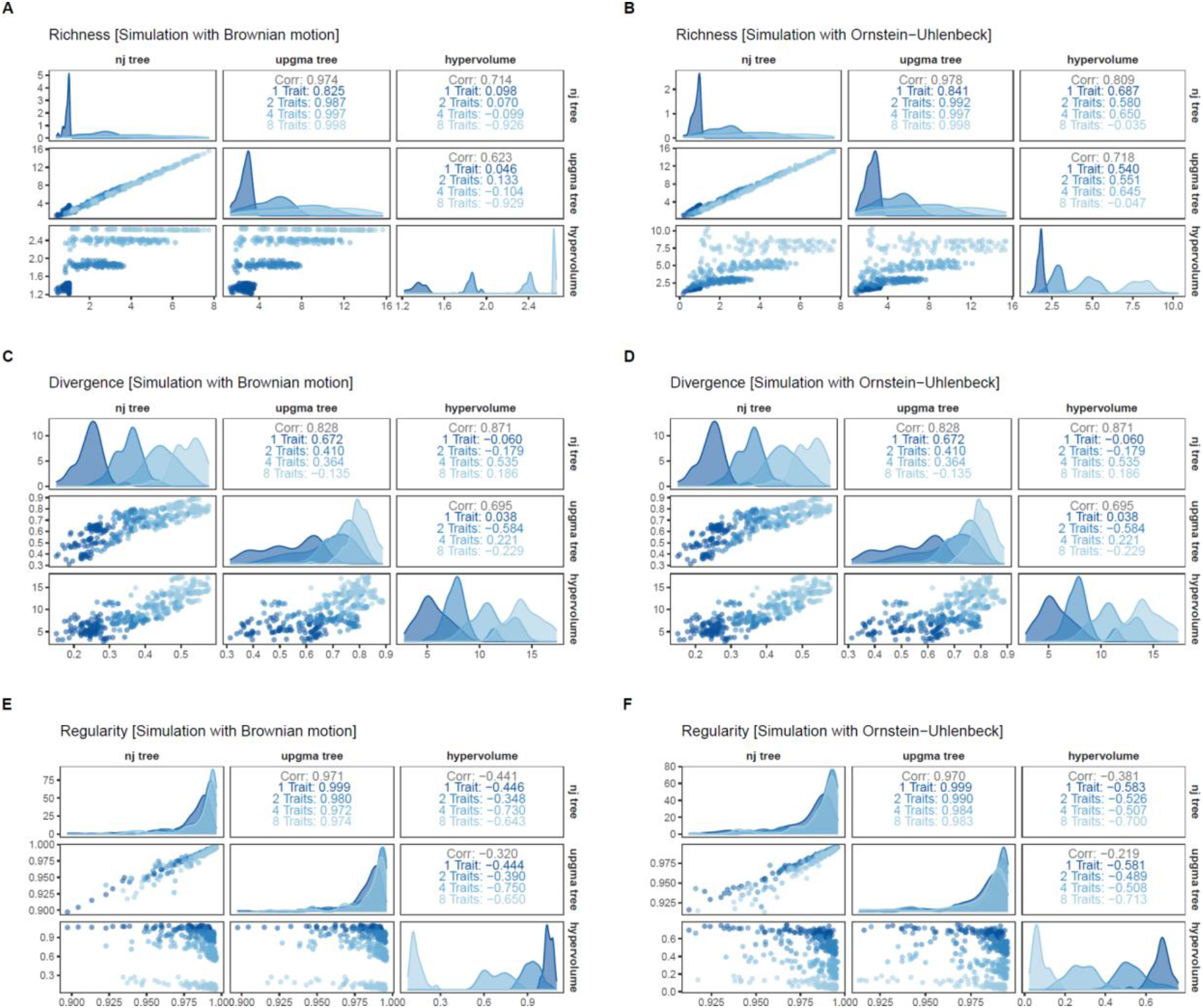
Pairwise Spearman’s correlations among estimations of functional richness (**A, B**), divergence (**C, D**), and regularity (**E, F**) with NJ trees, UPGMA trees, and kernel-density n-dimensional hypervolumes. For each panel plot, density plots on the diagonal display the distribution of values. Bivariate scatter plots are displayed below the diagonal and the correlation values above the diagonal.

The quality of the simulated functional spaces obtained by the NJ method was superior to those constructed by UPGMA, for all the combinations of number of species per community (from 10 to 100 species), number of traits per species (one, two, four and eight) and evolutionary processes used to generate the traits (BM and OU) (Appendix S1). It is worth mentioning that the quality of the functional spaces constructed by NJ for communities with only one trait was always 1. Results of the mixed model analysis confirmed that NJ performance, in terms of functional space quality, was significantly better than UPGMA (Table 1).

**Table 1.**
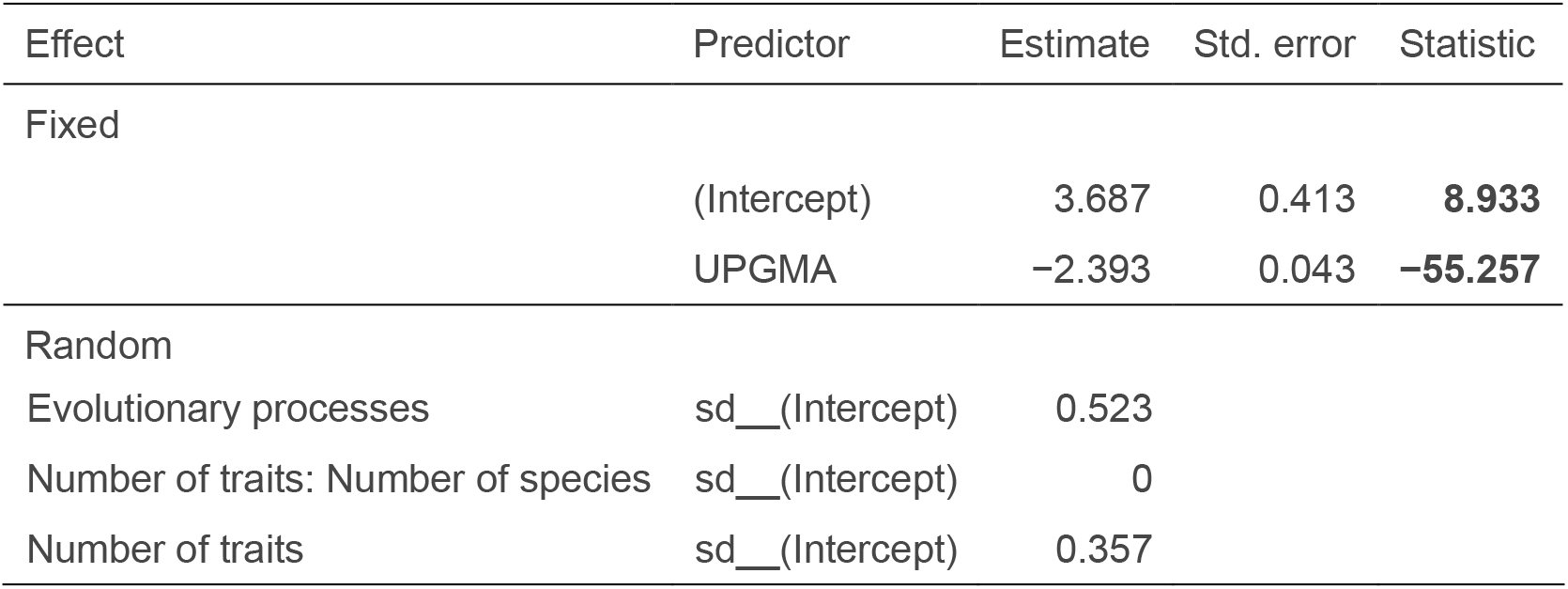
Summary of a mixed model for the quality of functional space, where we included method (NJ and UPGMA) as a fixed parameter predictor (Fixed) and allowed the intercept to vary (Random) across number of traits per species within blocks of number of species per community, and across evolutionary processes. Significant estimates are in bold.

| Effect | Predictor | Estimate | Std. error | Statistic |
| --- | --- | --- | --- | --- |
| Fixed | (Intercept) | 3.687 | 0.413 | **8.933** |
|  | UPGMA | -2.393 | 0.043 | **-55.257** |
| Random |  |  |  |  |

|  |  |  |
| --- | --- | --- |
| Evolutionary processes | sd__(Intercept) | 0.523 |
| Number of traits: Number of species | sd__(Intercept) | 0 |
| Number of traits | sd__(Intercept) | 0.357 |

The quality of the functional spaces built using the NJ method was similar to that obtained with four dimensions (half the maximum number of axes) using PCoA. The performance of NJ was higher than multidimensional spaces with two or three dimensions, but lower than multidimensional spaces with more than four dimensions (Table 2). A degree of caution is required when interpreting the mixed models results. Specifically, the variance estimates associated with the evolutionary processes (variable with 2 levels) and the number of traits (variable with 4 levels) should be regarded cautiously, due to the small number of levels involved.. Nevertheless, it must be emphasised that the main purpose of fitting the model was to compare the performance of methods in terms of functional space quality and not the effects of evolutionary processes and the number of traits per se.

**Table 2.**
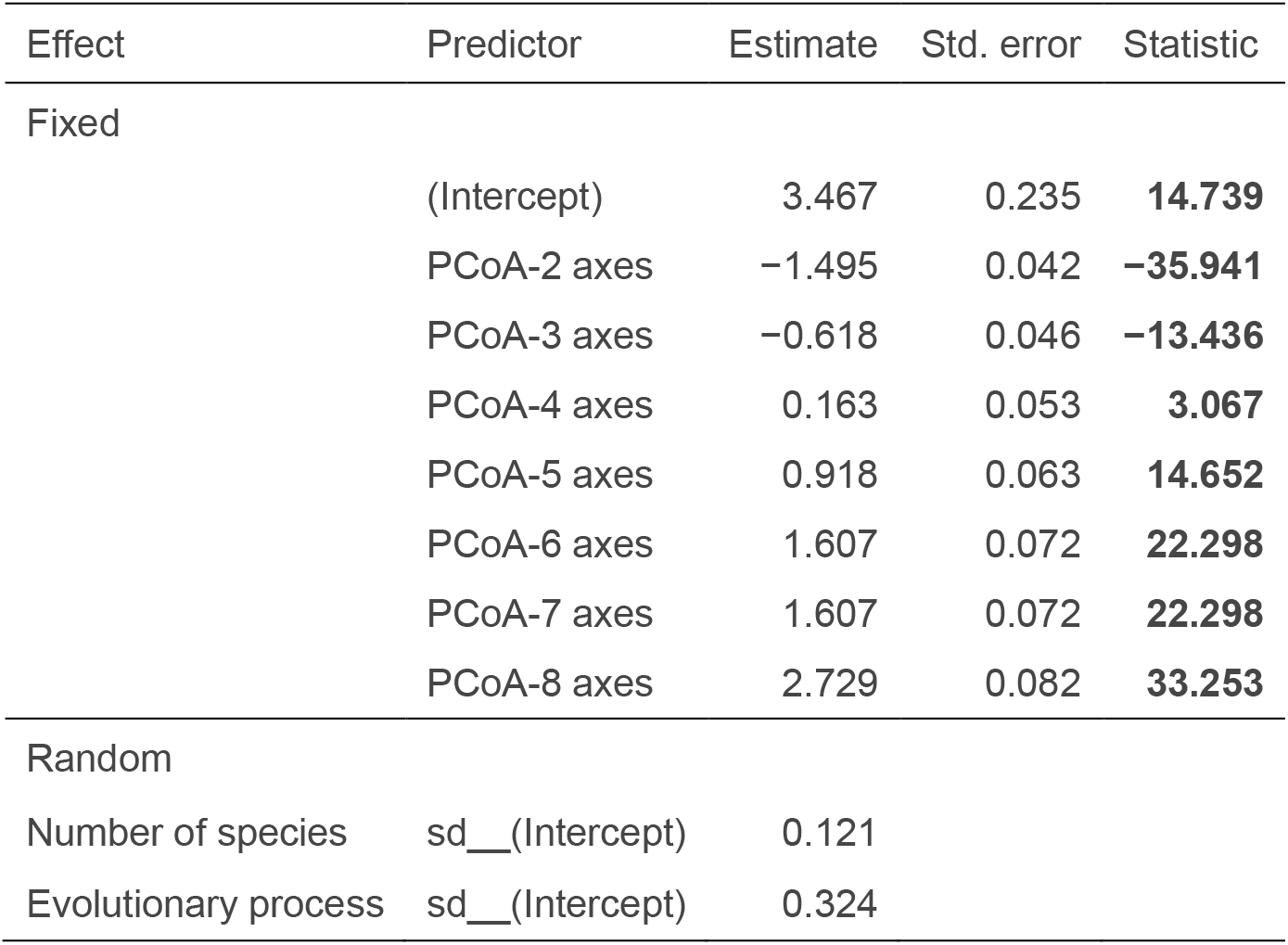
Summary of the mixed model for the quality of functional space, where we included method (NJ and from two to eight dimensions corresponding to the axes provided by a PCoA) as a fixed parameter predictor (Fixed) and allowed the intercept to vary (Random) across the number of species per community and evolutionary processes. Significant estimates in bold.

Regarding the sensitivity to outliers, NJ and UPGMA were found to be similar for all simulations regardless of the number of traits (Appendix S2). In contrast, hypervolumes were more sensitive, with higher differences between initial and final richness values after excluding the most unique species in each community for most scenarios.

In regard to the empirical test focused on the “kiwi problem”, the quality of the functional space for birds using NJ was 0.994, compared with 0.953 for UPGMA and 0.996 for two PCA axes (>0.999 for three or more axes). The exclusion of kiwis from the community, i.e., decreasing ~5% of the species richness, led to a decrease in functional richness of 10 and 14% for UPGMA and NJ respectively. In contrast, functional richness as measured using convex hulls was reduced by 73% and using kernel density hypervolumes by 42% (Fig. 4).

**Fig. 4.**
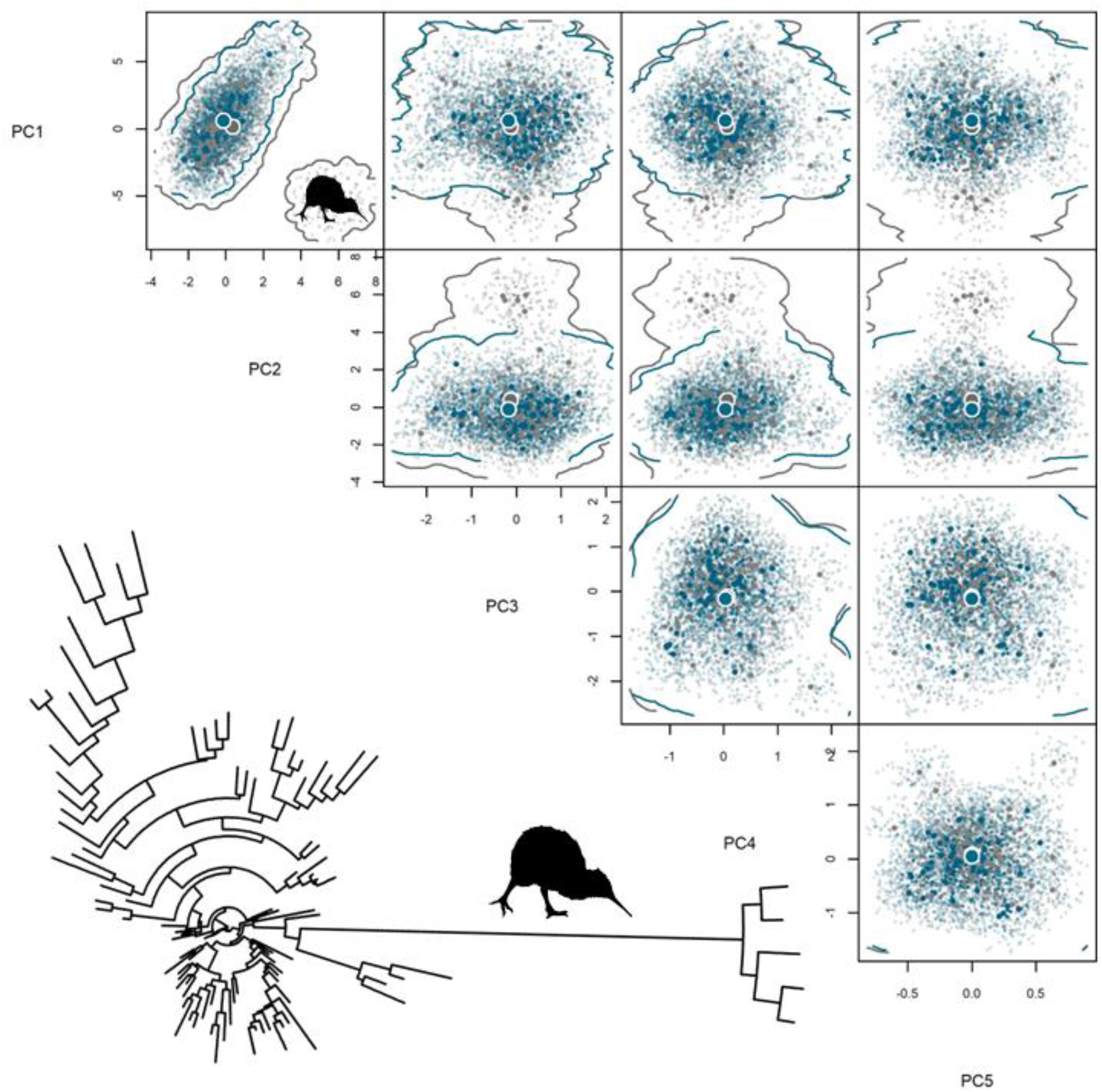
NJ functional tree with the edge leading to the five kiwi species highlighted by the silhouette, and the multidimensional space of studied birds with (grey) and without (blue) kiwis included. Five axes were selected for analyses, adding to 99% of explained variance. Note that, when excluding the kiwis, it is not just the space they occupy that is lost, but the space representing the remaining 100 species also “shrinks”, as the average distance between species decreases. For convex hulls the difference is even higher, as all the empty intervening space is also lost. Kiwi silhouette by Ferran Sayol.

## Discussion

The study of functional diversity is a burgeoning research area in ecology and evolution, with numerous methodological developments during the last couple of decades (Mammola et al., 2021; De Bello et al., 2021; Palacio et al., 2022). In contrast to the study of taxonomic or phylogenetic diversity, where the methodological approaches to quantify diversity are relatively established, there is still much discussion around how to best represent and measure FD across its dimensions, namely richness, divergence and regularity. Raw data, different tree representations, or representations based on multidimensional spaces, all have their strengths and weaknesses (Mammola et al., 2021). Here we propose a novel approach to measuring FD that combines many of the advantages of these different approaches, while minimising the disadvantages.

### Statistical properties

Our results indicate that the NJ method is more accurate than UPGMA (and similar methods such as minimum spanning trees) in representing the functional space occupied by a given community, i.e., the quality of the trait space. It is on par with multidimensional representations with up to four dimensions in simulated scenarios that cover a large variety of real-world situations. Given the “curse of dimensionality” (Bellman, 1957) of the hyperspace – the mathematical and computational difficulty of dealing with many dimensions simultaneously, and the implicit negative relationship between the number of dimensions and the volume of the hyperspace – a decrease in correlation between NJ trees and hypervolumes with increasing number of dimensions is expected. In general, NJ will be as accurate as hypervolumes in many situations and will present only small differences otherwise.

Multidimensional representations are known to have difficulties dealing with outliers, with substantially unique observations having disproportional effects on the quantification of FD. In the empirical example illustrating the “kiwi problem” (Fig. 4), excluding the kiwis from the hypervolume construction does not just result in the loss of the space they occupy, but the space representing the remaining 100 species also “shrinks”, as the average distance between species decreases.. For the very commonly used convex hulls, the loss is even more severe, as it includes all the intervening functional space (i.e., the space between the kiwis and all other birds where the convex hull extends out) that is in fact not occupied by any real bird species. NJ trees can circumvent the “kiwi problem” by generating a representation that is less sensitive to the large functional differences between kiwis and the remaining birds, but that is of higher quality than UPGMA trees.

### Comparing different facets of diversity

As with UPGMA and other tree methods, taxonomic diversity can be represented as a star-like NJ tree, and in fact the construction of a NJ tree starts with a starlike tree. This means that TD and FD are comparable using the same methods, although for TD they are usually simplified for speed and ease of use.

Crucially, we demonstrate that hyperspatial representations are not comparable with tree representations that are often used for quantifying PD. As seen in Fig. 2, even for the same data, trees and hypervolume values of richness, divergence or regularity have little to no correlation. This implies that, if one uses phylogenetic trees to measure PD and hypervolumes to measure FD, any differences in patterns will be due to both differences in community composition and the mathematical properties of the indices, with no possibility to disentangle these two effects. We should note that trees used for quantifying PD can be used with any method, including NJ, Bayesian or any other that results in a tree (ultrametric or not, dated or undated). The mathematics will be similar and hence comparability is warranted.

### Ease of use

As with other tree representations, NJ works directly with distances between species. The choice of distance is critical, although Gower’s distance is often preferred as it allows for the use of continuous, ordinal, binary and categorical variables (Pavoine et al., 2009). When only continuous variables are used, as in our empirical example, Euclidean distances are generally preferred. In any case, this decision is almost always simpler to make than the ones involved in the use of multidimensional methods, which include the number of axes to use, which method to use for estimating the kernel density, and the many parameters that can influence the results in substantial ways when building more complex representations.

The use of certain distance measures, such as Gower’s distance, allow for missing trait values, with no need for imputation. In addition, some of the methods for building NJ trees allow for missing distances between pairs of species (Criscuolo & Gascuel, 2008). The flexibility provided in both these steps, calculating distances and building trees, will help circumvent the many gaps that most trait databases have, particularly for taxa less well studied than birds (e.g., Pekar et al. 2022; Shirey et al. 2022).

The construction of NJ trees is extremely fast, orders of magnitude faster than hypervolumes, which can be an advantage for large datasets or simulations or null models requiring many repeated calculations. In addition, it is possible to at least visually estimate to a close approximation many of the metrics derived from tree-like representations (e.g., richness), a task that is much harder for multidimensional representations. This helps avoid errors in data input and/or coding, as many major errors will be obvious through inspection of the tree plot.

### Caveats

The main caveat of using NJ is the lack of apparent connection between trees and the intuitive representation of the Hutchinsonian niche (Mammola et al., 2021). It is intuitive to imagine the functional space occupied by a group of species as a multidimensional concept depicting its many functional dimensions. Conversely, the connections between species in a tree are not natural in the sense that they do not represent real connections in the community, only the closest path between them in the tree.

A second caveat is a potential lower flexibility than probabilistic hypervolumes to consider the abundances of species in the different metrics. The trait space is largely homogeneous in the way it is occupied, although abundances could theoretically be represented by the density of connections in parts of the tree. In addition, if intraspecific data are available, one can build trees using individuals instead of species, by-passing this issue. Intraspecific trait data are increasingly seen as being crucial to understanding how organisms interact (Tautenhahn et al., 2019; He et al., 2021; Wong & Carmona, 2021). Given that intraspecific trait data are not always available at the community level, one workaround is to simulate intraspecific variability from compound measures such as the standard deviation of a given trait, which could approximate the kernel-density approach using trees.

## Conclusions

In this study, we have proposed a novel approach to representing functional space and calculating FD that enables the quantification of its different dimensions in ways that combine the strengths of previously proposed FD frameworks. Extensive research on the properties of phylogenetic trees has been undertaken, and, using the NJ framework presented here, in the future these advances can be used to study different properties of ecological systems using functional trees (e.g., Ning et al., 2020). The mathematics underpinning NJ are already extensively developed, and thus the use of NJ trees opens up the possibility of testing new hypotheses for FD in the same way as has been done for PD. By combining ease and speed of use, low distortion of functional space, low sensitivity to outliers, and comparability with PD measures, the use of NJ seems a promising approach for the future. More broadly, there are other methods available for building trees that are not considered here, but that could also provide new and advantageous ways to represent functional diversity (e.g., Wheeler, 2021). As such, we would argue that further exploration and testing of alternatives to the commonly used functional tree construction approaches (e.g., UPGMA) will likely prove rewarding in the study of functional diversity going forward.

## Data Availability

All data are available through the AVONET database (Tobias et al., 2022).

## Author contributions

Pedro Cardoso and Jose Carlos Carvalho conceived the ideas and led the writing of the manuscript. Pedro Cardoso, Thomas Guillerme, Stefano Mammola, Thomas J Matthews, Caio Graco-Roza and Jose Carlos Carvalho designed methodology and analysed the data. All authors contributed critically to the drafts and gave final approval for publication.

## APPENDIX S1

**Fig. S1.1.**
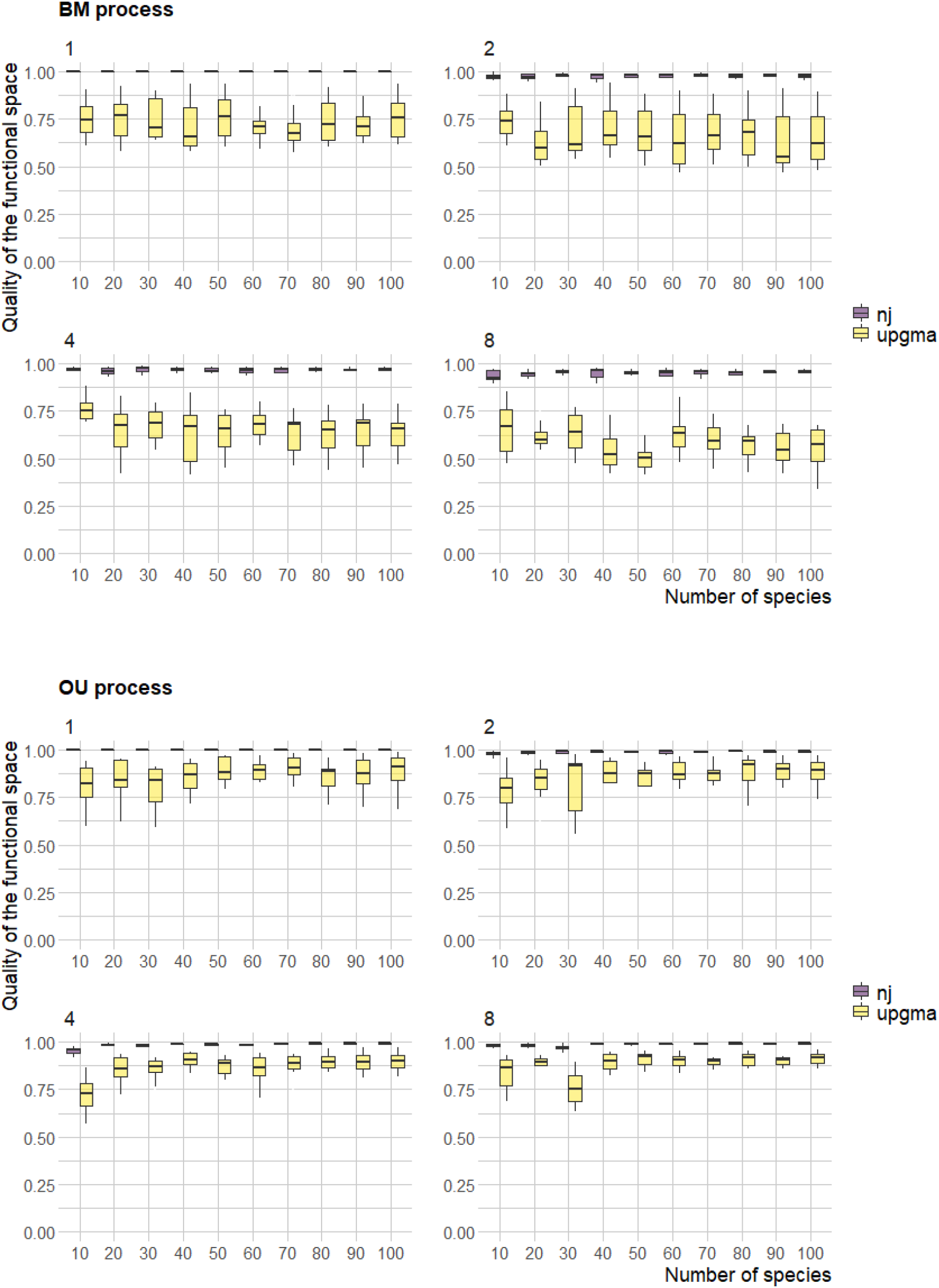
Comparison of the quality of the functional spaces obtained by NJ and UPGMA methods across different sets of simulations. Boxplots show the distribution of the functional space quality in 10 simulations for each combination of number of species per community (from 10 to 100 species), number of traits per species (1, 2, 4 and 8) and evolutionary processes used to generate the traits (BM and OU).

**Fig. S1.2.**
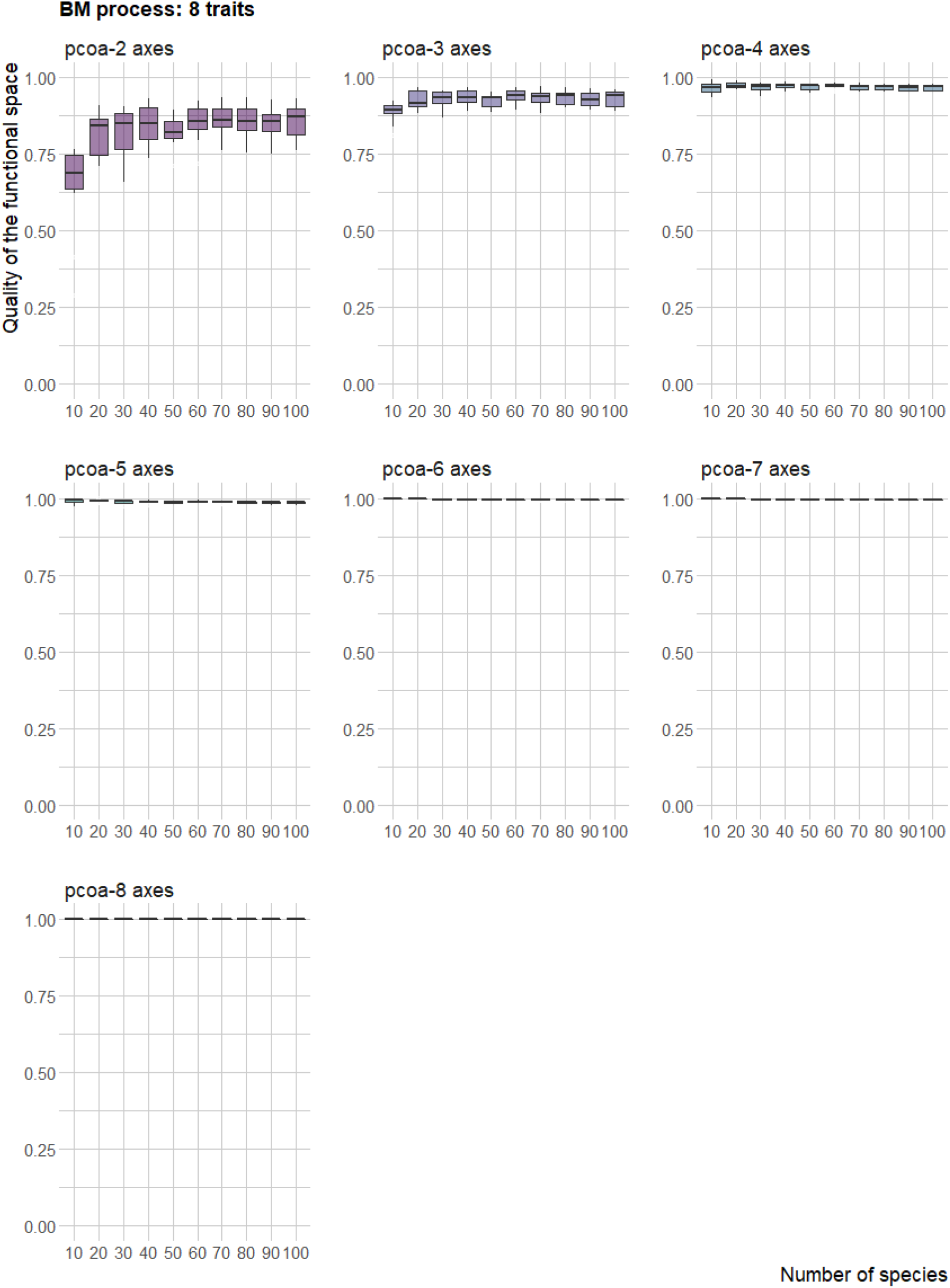
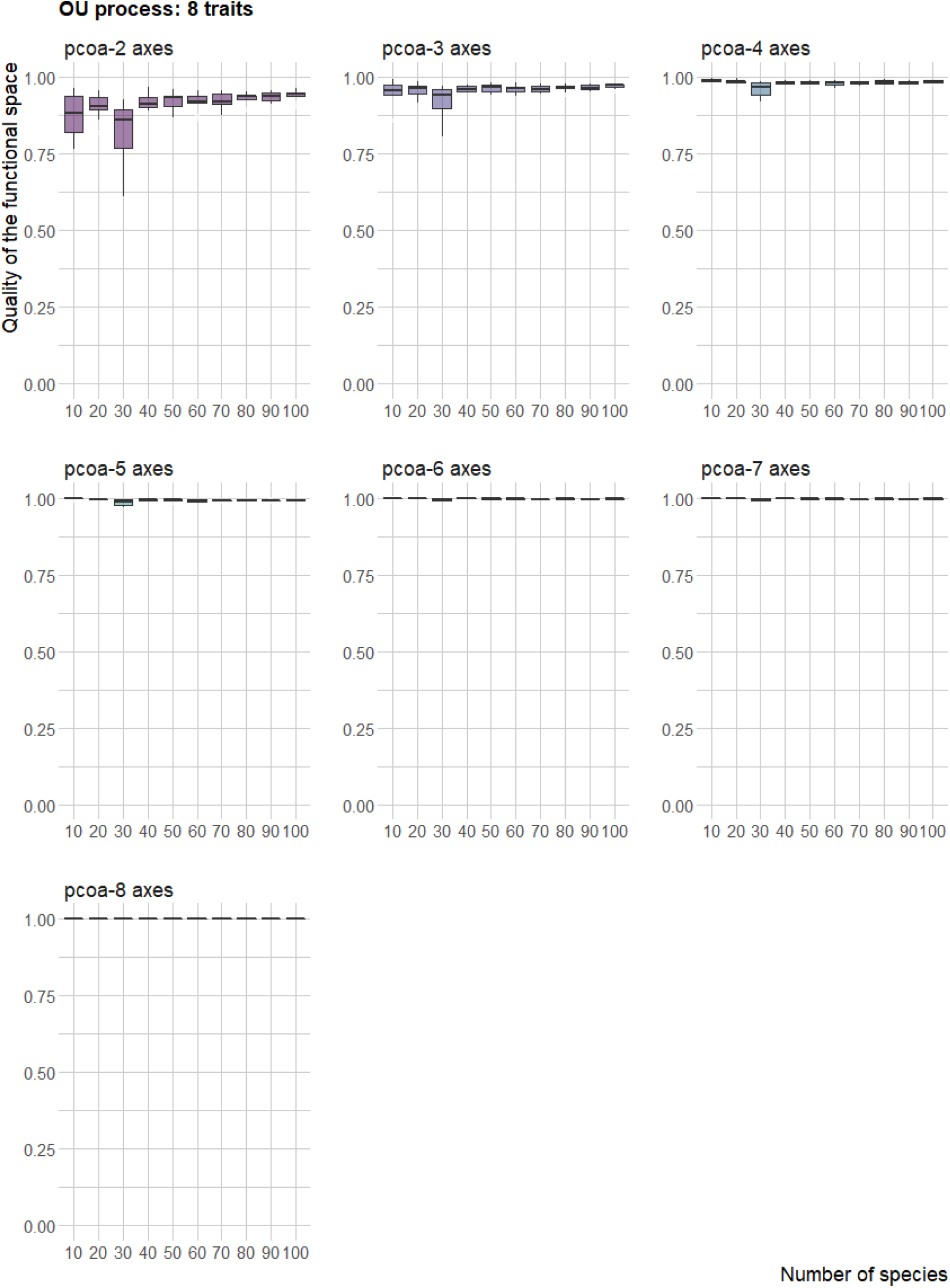
Quality of the multidimensional functional spaces (from two to eight dimensions) corresponding to the axes provided by a PCOA. Boxplots show the distribution of the functional space quality in 10 simulations with different number of species per community (from 10 to 100 species), using 8 traits generated with two evolutionary processes (BM and OU).

## APPENDIX S2

**Fig. S2.1.**
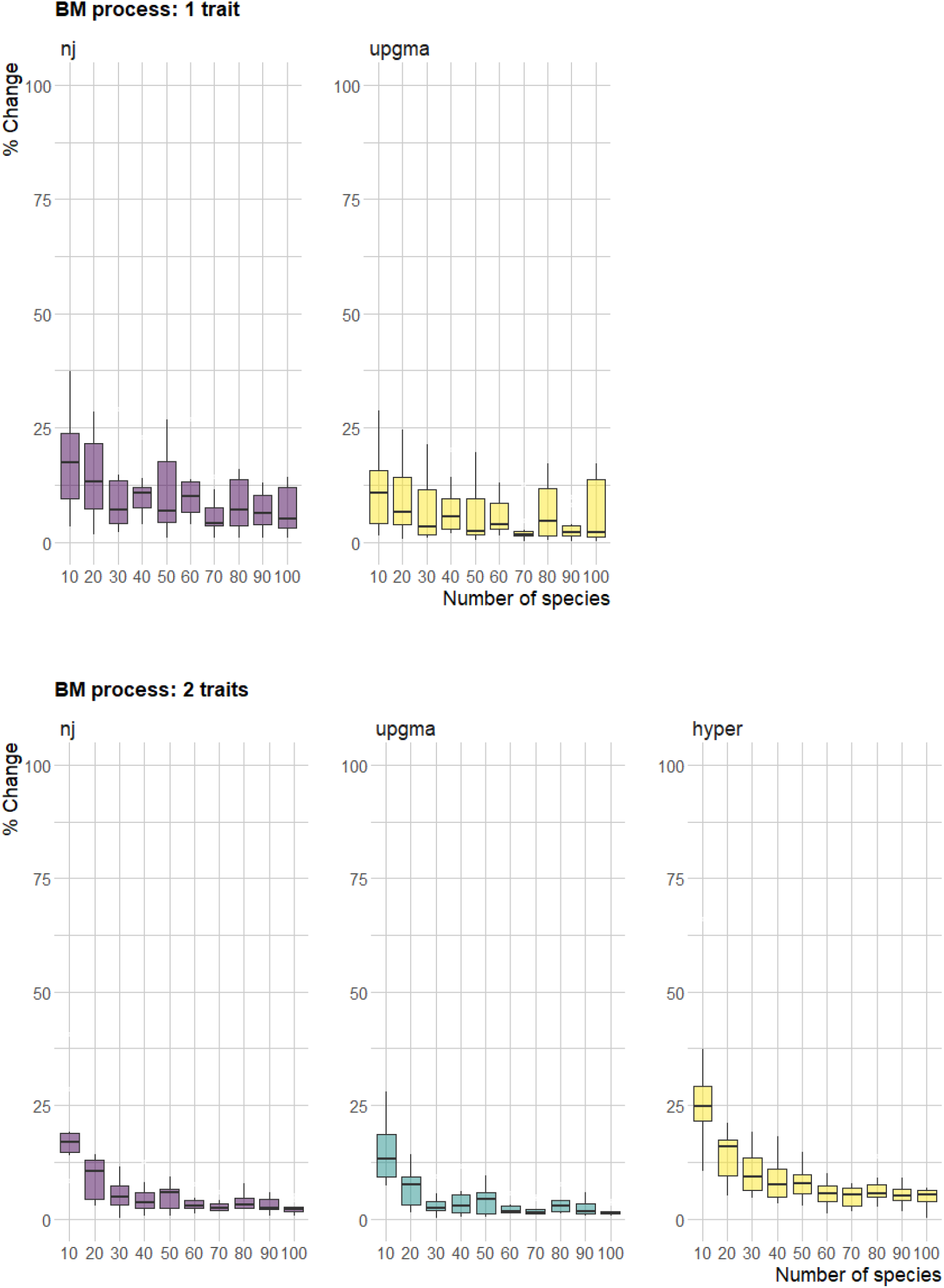
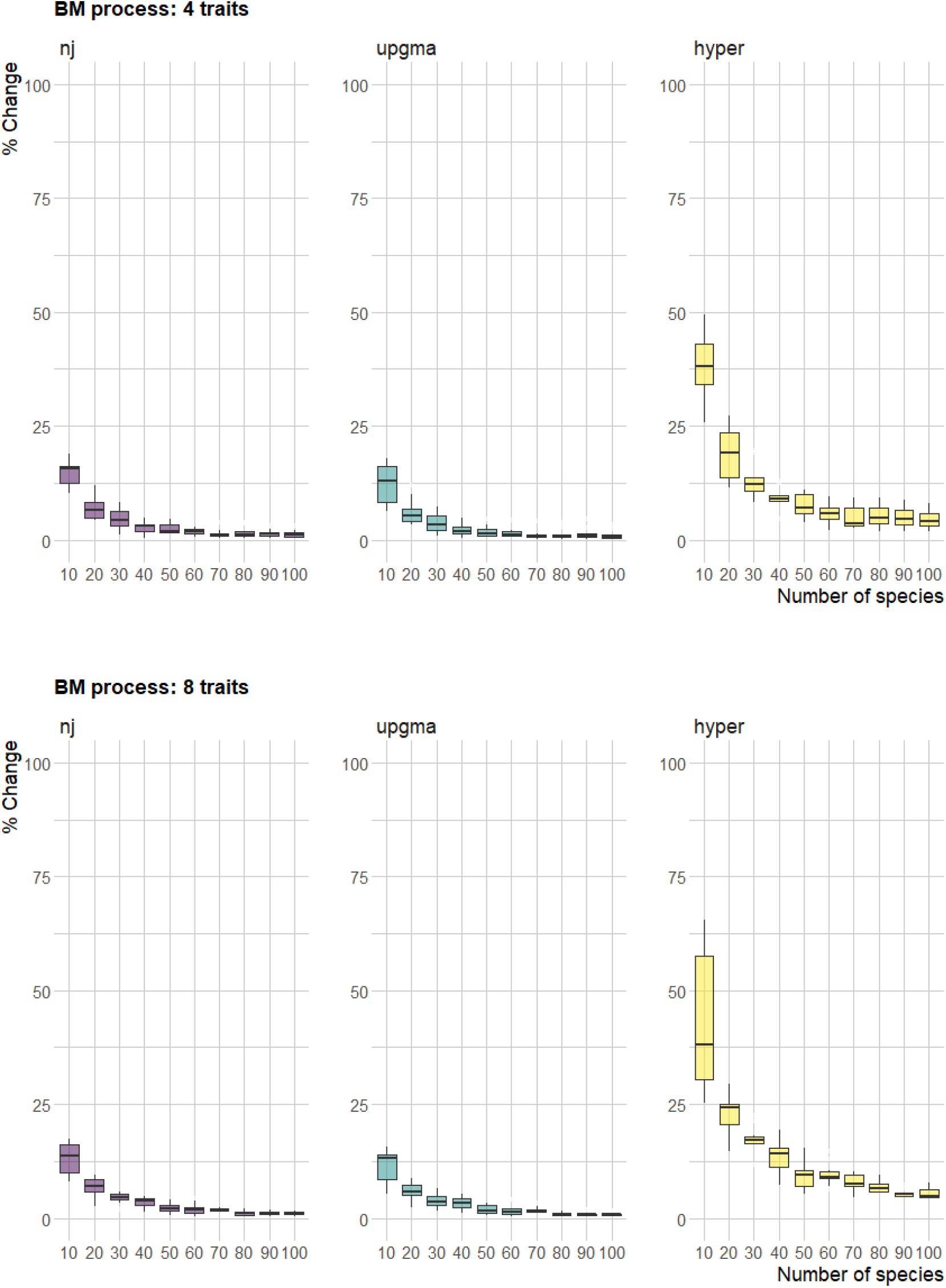
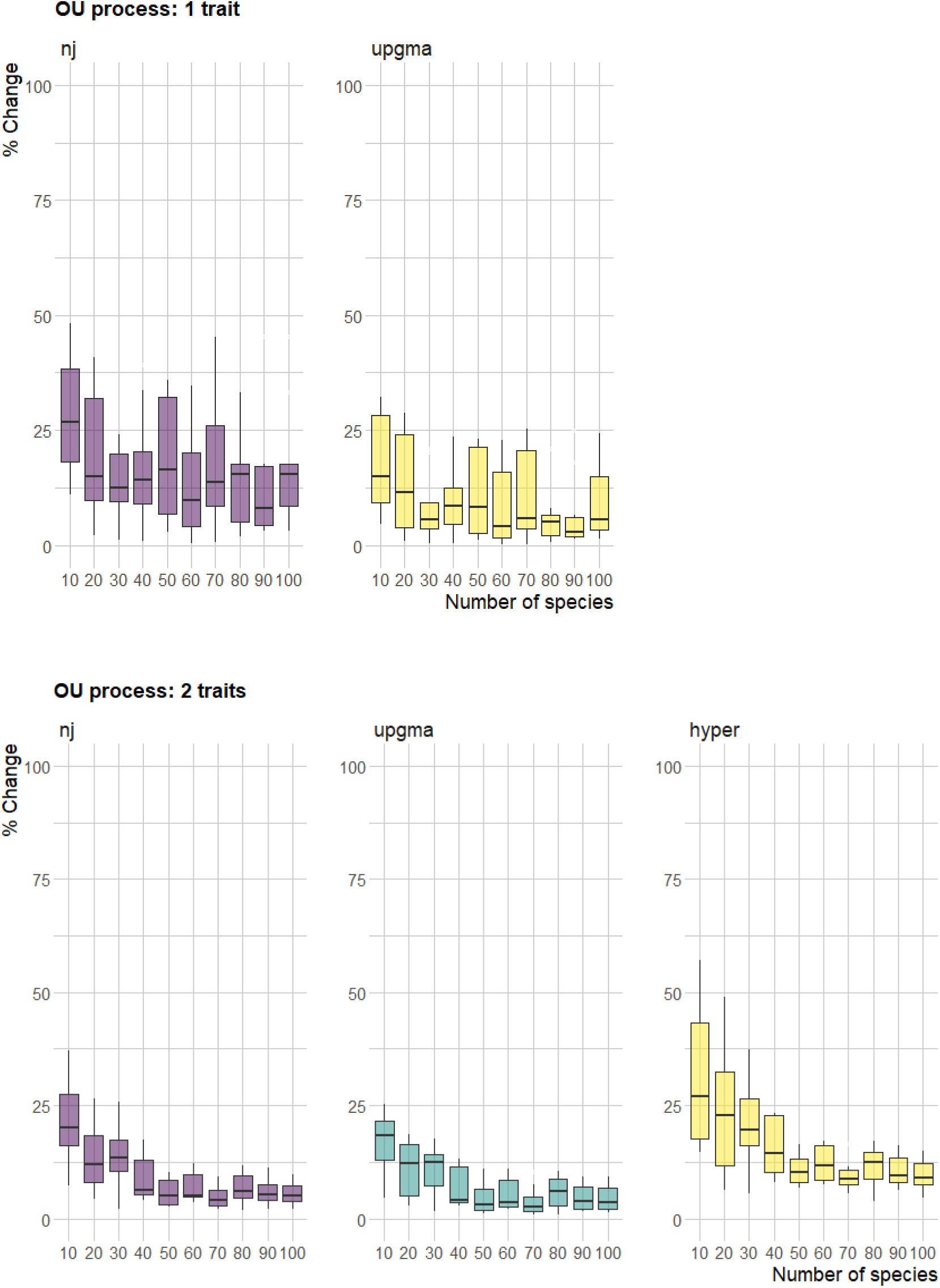
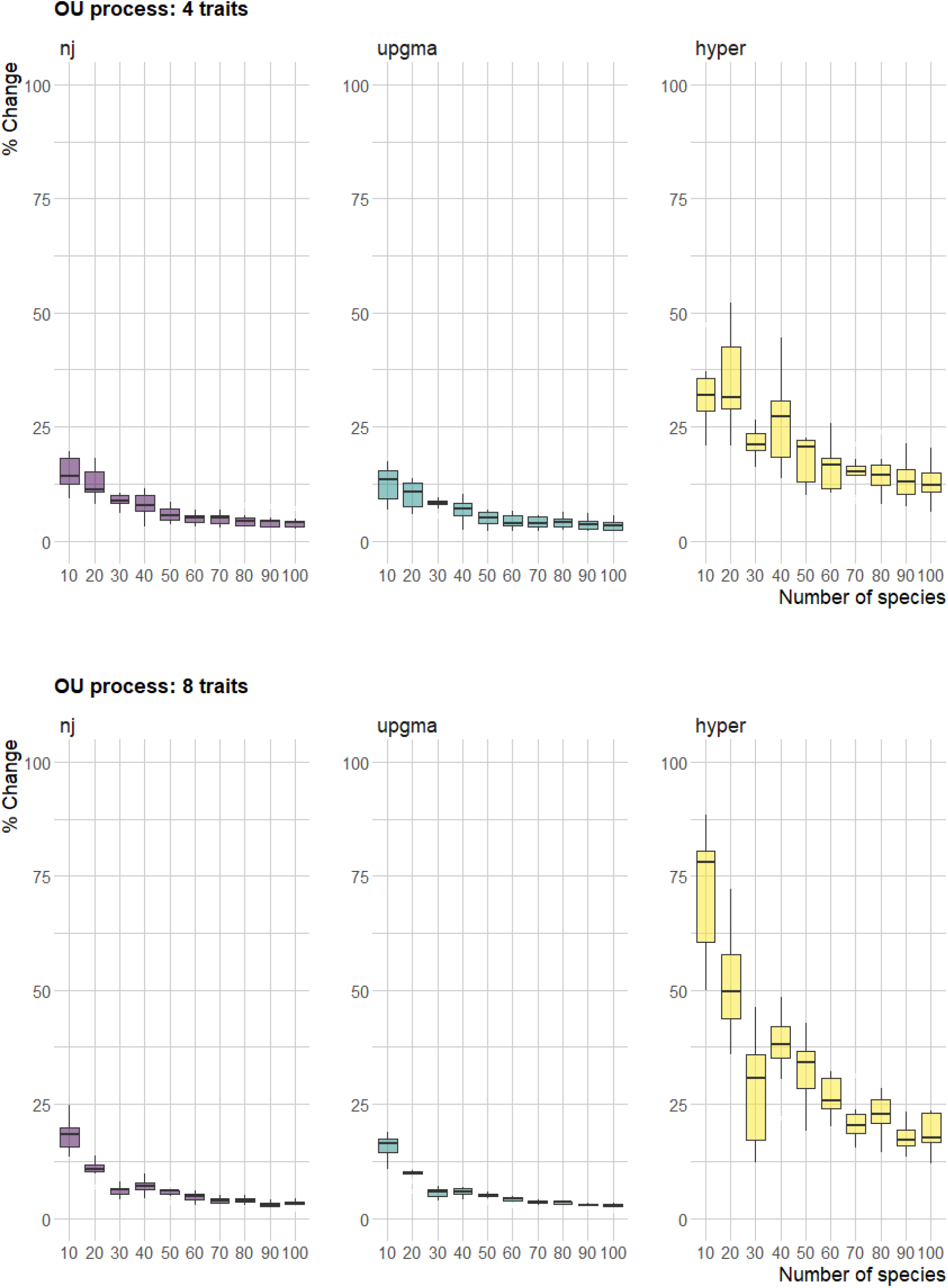
Sensitivity to outliers of NJ, UPGMA and kernel-density hypervolumes methods across different sets of simulations. Sensitivity is expressed as the percentage of change in the values of richness, before and after deleting the species with higher uniqueness (outlier) in each community. Boxplots show the distribution of sensitivity values in 10 simulations for each combination of number of species per community (from 10 to 100 species), number of traits per species (1, 2, 4 and 8) and evolutionary processes used to generate the traits (BM and OU). Note that for sets with only one trait only NJ and UPGMA were calculated.

